# Single-unit activations confer inductive biases for emergent circuit solutions to cognitive tasks

**DOI:** 10.1101/2024.11.23.625012

**Authors:** Pavel Tolmachev, Tatiana A. Engel

## Abstract

Trained recurrent neural networks (RNNs) have become the leading framework for modeling neural dynamics in the brain, owing to their capacity to mimic how population-level computations arise from interactions among many units with heterogeneous responses. RNN units are commonly modeled using various nonlinear activation functions, assuming these architectural differences do not affect emerging task solutions. Contrary to this view, we show that single-unit activation functions confer inductive biases that influence the geometry of neural population trajectories, single-unit selectivity, and fixed point configurations. Using a model distillation approach, we find that differences in neural representations and dynamics reflect qualitatively distinct circuit solutions to cognitive tasks emerging in RNNs with different activation functions, leading to disparate generalization behavior on out-of-distribution inputs. Our results show that seemingly minor architectural differences provide strong inductive biases for task solutions, raising a question about which RNN architectures better align with mechanisms of task execution in biological networks.

---

Continuous-time recurrent neural networks (RNNs) offer a versatile framework for modeling mechanisms of cognitive computations in the brain^1–3^. Similar to biological neural circuits, RNNs consist of many interconnected units with nonlinear activation function, which mimics the nonlinear input-to-output transformation performed by either an individual neuron or an entire group. Whether through training to perform behavioral tasks^4–6^ or by directly fitting recorded neural activity^7–9^, RNN units develop heterogeneous responses similar to the mixed selectivity observed in brain recordings^2^. Thus, RNNs serve as computationally tractable models that capture key features of biological neural networks. Reverse-engineering task solutions emerging in RNNs through training provides hypotheses for how biological networks may execute similar cognitive tasks^1, 5, 8, 10^.

Continuous-time RNNs are universal approximators of any dynamical system. Specifically, RNNs can approximate the desired dynamics with arbitrary precision in a subset of output units, provided a sufficient number of hidden units in the network^11, 12^. Although the proof of this result holds for smooth and bounded sigmoid-like activation functions, the empirical evidence suggests that RNNs with rectified linear (ReLU) activation may approximate complex dynamics as well^3, 6^. Accordingly, it is commonly assumed that a specific choice of activation function is inconsequential to the mechanisms that emerge in RNNs, as long as the networks can be adequately trained to perform the task. Supporting this assumption, a comprehensive study of RNNs with different architectures found that, despite some differences in the geometry of neural dynamics, they use similar computational scaffolds, as characterized by the topological structure of fixed points^13^. This premise led to numerous studies employing a variety of activation functions to model biological networks, including sigmoid^14, 15^, ReLU^3, 6, 10, 16, 17^, and hyperbolic tangent (tanh)^4, 5, 18–21^, with the latter being the most common choice. However, whether these architectural choices are truly inconsequential to the circuit mechanisms emerging in RNNs through training has not been systematically tested.

We hypothesized that seemingly minute differences in the geometry of neural representations across RNNs with different architecture^13, 22^ may reflect deeper distinctions in the underlying circuit mechanisms driving behavior. To test this hypothesis, we analyzed RNNs with six distinct architectures trained on a range of behavioral tasks. We used three common activation functions (ReLU, sigmoid, and tanh) and, for each, trained RNNs with and without the Dale’s law constraint on the connectivity (restricting units to be either excitatory or inhibitory). Across all RNNs, we compared the geometry of neural population trajectories, single-unit selectivity, and fixed point configurations for all task conditions. All these attributes differed across RNNs with varying activation functions, with tanh networks diverging the most from both sigmoid and ReLU RNNs. Using a model distillation approach that fits RNN responses and task behavior with a low-dimensional latent circuit^10^, we uncovered that these differences in neural representations indeed arose from distinct circuit solutions used by the RNNs to solve the same task. Moreover, these circuit solutions made disparate predictions for how RNNs would respond to out-of-distribution inputs, which were definitively confirmed through simulation tests.

Our results reveal that single-unit activation functions confer inductive biases for circuit solutions that emerge in RNNs through training on cognitive tasks. These distinct circuit solutions are mirrored in neural representations, single-unit selectivity, and fixed point configurations, and produce disparate behavioral outcomes for out-of-distribution inputs. Our findings imply that conclusions about mechanisms of task execution derived from reverse-engineering RNNs may depend on subtle architectural differences, emphasizing the need to identify architectures with inductive biases that most closely align with biological data.

## Results

How profound are differences across networks trained with varying architectures, such as differing activation functions and connectivity constraints? To answer this question, we trained RNNs with various architectures on a range of behavioral tasks. We used three common activation functions (ReLU, sigmoid, tanh) and, for each, trained 100 networks both with and without the Dale’s law constraint on the connectivity (Dale, No Dale), resulting in six distinct architectures. All RNNs were trained on the same task inputs and outputs to a similar performance level (Table 1). We then selected 30 top performing networks from each architectural class for the analysis.

**Table 1.** Coefficient of determination *r*^2^ between the RNN output and the target. Data are mean± std across top 30 RNNs for each task and architecture.

|  | ReLU |  | Sigmoid |  | Tanh |  |
| --- | --- | --- | --- | --- | --- | --- |
|  | Dale | No Dale | Dale | No Dale | Dale | No Dale |
| **CDDM** | $92 \pm 0.3\%$ | $93 \pm 0.6\%$ | $90 \pm 1.2$ | $87 \pm 1.5\%$ | $88 \pm 0.4\%$ | $88 \pm 0.5\%$ |
| **Go/NoGo** | $95 \pm 0.6\%$ | $95 \pm 0.6\%$ | $95 \pm 0.8\%$ | $95 \pm 0.7\%$ | $95 \pm 0.8\%$ | $95 \pm 0.7\%$ |
| **Memory Number** | $99 \pm 0.1\%$ | $99 \pm 0.1\%$ | $98 \pm 0.1\%$ | $99 \pm 0.2\%$ | $99 \pm 0.1\%$ | $99 \pm 0.1\%$ |

Across all RNNs, we compared neural representations using population trajectories and single-unit selectivity, and dynamical mechanisms characterized by the configurations of fixed points and trajectory end points. We further extracted circuit mechanisms driving task behavior in these RNNs and tested their generalization performance on out-of-distribution inputs. We first present our findings in detail focusing on a context-dependent decision-making task (Fig. 1, Methods: Context-dependent decision-making task), and then show that these observations generalize to other tasks as well.

**Figure 1.**
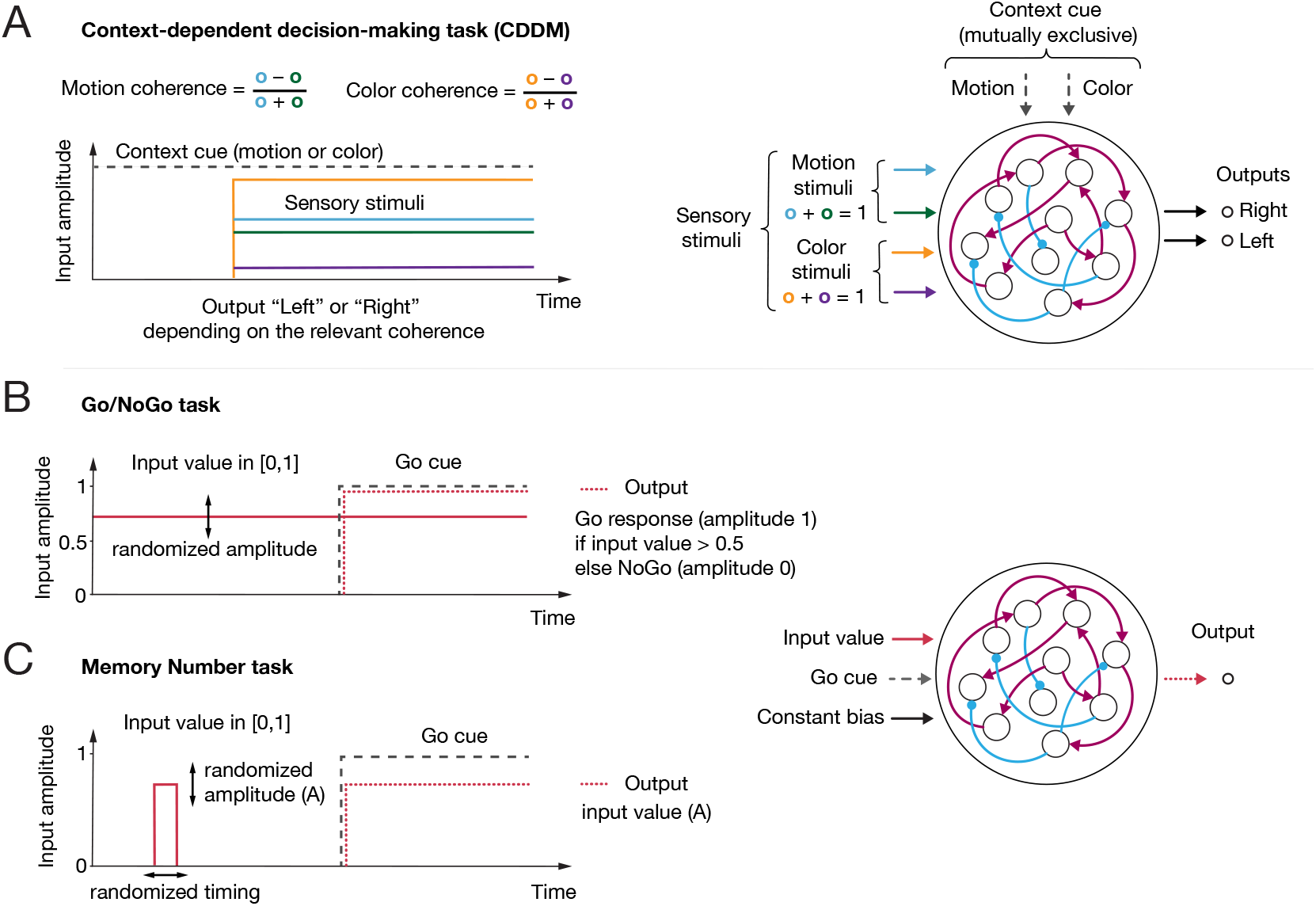
Tasks used for RNN training. **A**. Context-dependent decision-making (CDDM) task (right). RNN receives two modalities of sensory inputs, termed ‘Motion’ and ‘Color’. For each modality, two positively-constrained input channels provide momentary evidence for left and right choice, respectively. The difference between the mean right and left input defines stimulus coherence, with values ranging from ™1 to 1. Two context channels supply the cued context input, with only one context channel active on each trial indicating the relevant sensory modality. The network is required to output ‘Left’ or ‘Right’ decision on the corresponding output channel based on the signed coherence of the relevant sensory input. Time-course of the CDDM task (left). The ‘Context cue’ is present throughout the trial, indicating either ‘Motion’ or ‘Color’ context. Following the context cue, four channels convey sensory stimuli (2 sensory modalities with 2 channels per modality). The network is required to produce an output as soon as the sensory inputs are supplied. **B, C**. Go/NoGo and MemoryNumber tasks have shared input and output structure (right). **B**. Go/NoGo task. A single input value chosen from (0, 1) range is presented throughout the trial. Upon presentation of the ‘Go cue’, the network is required to output ‘Go’ response with amplitude 1 if the input value is above 0.5 threshold, and produce ‘No Go’ response with amplitude 0 if the input value is below 0.5. **C**. Memory Number task. An input value with randomized amplitude (A) is briefly presented at the beginning of the trial within a randomize time window. Upon receiving a ‘Go cue’, the RNN is required to output the same value A.

### Differences in representations across RNN architectures

We compared neural representations in trained RNNs by analyzing the geometry of trajectories in the neural population state space and tuning of single units in the selectivity space. Population trajectories and single-unit selectivity provide dual perspectives on the same neural representations^23, 24^. Consider a neural response matrix containing the activity of a single RNN with *N* units across *K* trials. This matrix has dimensionality of (*N, T* × *K*), where *T* is the number of time steps in a trial. The columns of this matrix define the neural population state space, where each axis represents activity of one unit and each point is a sample of population activity. The rows of the same matrix define the selectivity space, where each axis represents a response feature and each point is a neuron. Accordingly, we perform dimensionality reduction with Principle Component Analysis (PCA) on either the columns or rows of the neural response matrix to obtain the projected configurations of population trajectories and single-unit selectivity. To analyze population trajectories, we reduce the first dimension from *N* to *n*_PC_ and obtain a matrix with dimensionality (*n*_PC_, *T* ×*K*) containing a set of projected trajectories (Methods: Analysis of population trajectories). To analyze single-unit selectivity, we reduce the second dimension from *T* × *K* to *n*_PC_ and obtain a matrix with dimensionality (*N, n*_PC_) containing task tuning information for all RNN units (Methods: Analysis of single-unit selectivity).

For initial assessment, we visualized population trajectories of example RNNs by projecting them onto the first two PCs. The trajectories of ReLU and sigmoid RNNs are visually distinct from those of tanh networks (Fig. 2A). ReLU and sigmoid RNNs typically form symmetric, butterfly-shaped trajectory sets: the trajectories stay near the origin during the presentation of the context cue at the beginning of the trial, and gradually separate later in a trial when the sensory inputs are introduced. In contrast, the trajectories of tanh RNNs promptly diverge at the beginning of the trial, driven solely by the context inputs. Later in the trial, these trajectories further split according to the sensory inputs forming two sheets orthogonal to the context axis. Dale’s law connectivity constraint did not affect the geometry of population trajectories of tanh RNNs. In ReLU and sigmoid RNNs, however, the Dale’s law constraint led to more structured representations, with trajectories clustering according to the context and choice, whereas the trajectories varied more continuously in ReLU and sigmoid RNNs without Dale’s law.

**Figure 2.**
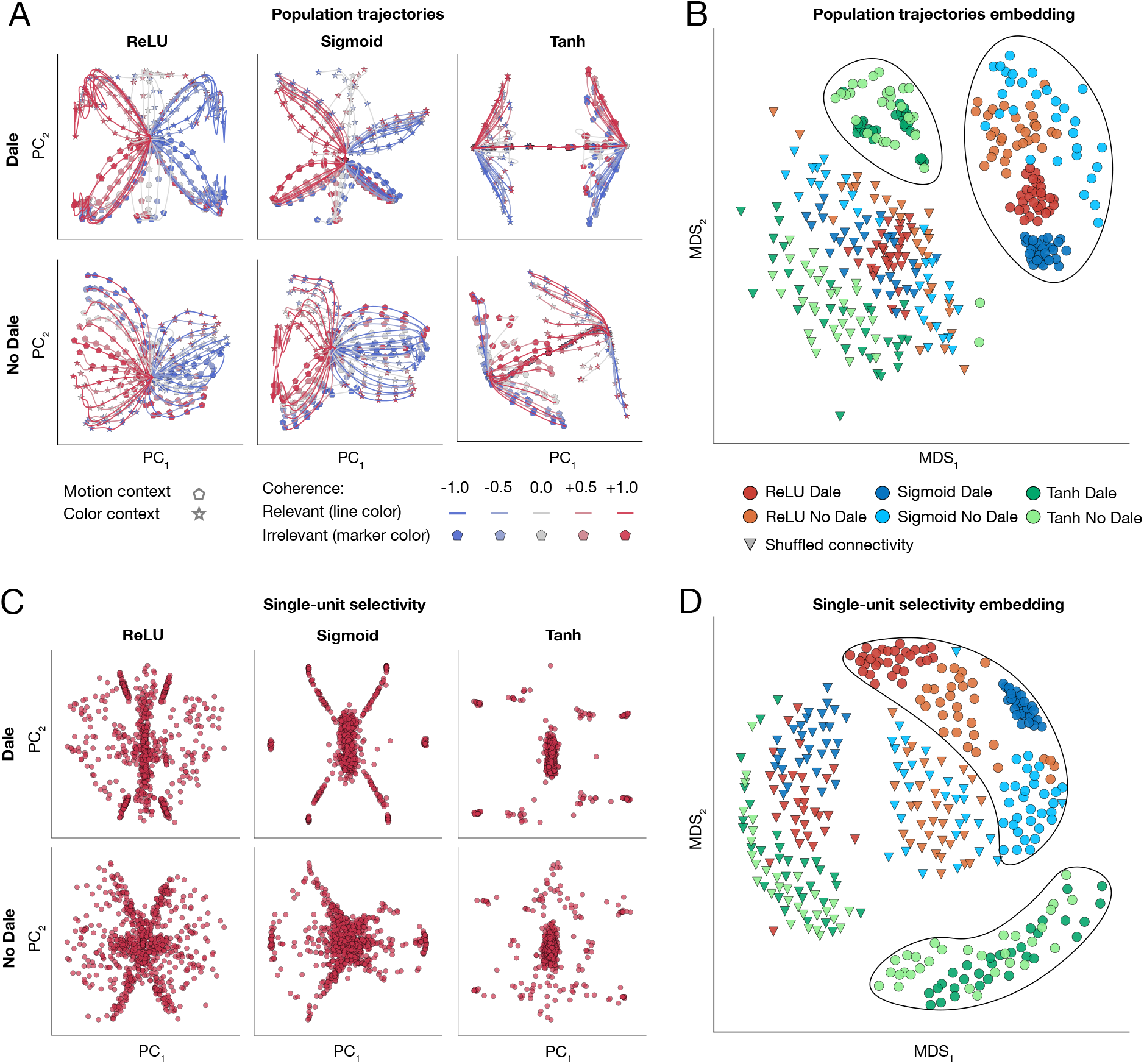
Comparison of population trajectories and single-unit selectivity across six RNN architectures trained on CDDM task. **A**. Population trajectories visualized by projecting onto the first two principal components (PCs) in example RNNs with different activation functions (columns) and connectivity constraints (rows). **B**. MDS embedding of population trajectory sets across all RNNs. Each point represents set of trajectories of a single RNN. ReLU and sigmoid networks form clusters distinct from tanh networks. Triangles represent trajectories of the same RNNs with shuffled connectivity matrices, used as a control. **C**. Single-unit selectivity visualized by projecting onto the first two PCs in RNNs with different activation functions (columns) and connectivity constraints (rows). Each point represents one unit. Each plot aggregates units from the top 30 RNNs, showing only units with activity levels above the 50^th^ percentile. **D**. MDS embedding of single-unit selectivity configurations across all RNNs. Each point represents one RNN. RNNs with each architecture form distinct clusters, with the tanh RNN cluster positioned further away from the others. In B and D, each RNN class is represented by top 30 RNNs (360 RNNs in total, including controls).

To systematically quantify these noticeable differences in trajectories across all RNNs, we embedded individual trajectory sets into shared two-dimensional space, where trajectory sets of each RNN are represented by a single point, and distances between points characterize similarity of trajectory sets across RNNs (Fig. 2B, Methods: Analysis of population trajectories). To obtain this embedding, we computed a pairwise distance metric as the mean squared error (MSE) between projected trajectories (*n*_PC_ = 10) of two RNNs after their optimal alignment with orthogonal Procrustes. We then embedded the distances between all RNN pairs in a two-dimensional space using multidimensional scaling (MDS)^13^, which aims to minimally distort all pairwise distances. This systematic analysis confirmed our initial observations from visualizing trajectory sets in example RNNs. The networks with different architectures form distinct clusters (Fig. 2B). Tanh RNNs with and without Dale’s connectivity constraint form a single cluster that is distinctly separated from clusters formed by ReLU and sigmoid RNNs. The clusters formed by ReLU and sigmoid RNNs without Dale’s connectivity constraint show higher spread than their constrained counterparts, indicating larger heterogeneity in population trajectories across networks.

We further examined single-unit selectivity configurations, which also differed across RNNs with different activation functions. Visualizing single-unit selectivity in example RNNs reveals striking differences between ReLU and sigmoid versus tanh networks (Fig. 2C). ReLU and sigmoid RNNs produce a cross-shaped pattern with continuously populated arms extending outward, whereas tanh RNNs display a large central cluster with a few distant, outlying units. We then computed pairwise distances between the single-unit selectivity configurations across all RNNs and embedded them into two-dimensional space using MDS. We computed a pairwise distance metric as MSE between two RNNs after matching their units in the selectivity space using iterative closest point (ICP) registration algorithm, which allows for one-to-many unit matching (Methods: Iterative closest point registration). The systematic analysis of single-unit selectivity configurations confirmed that tanh RNNs are distinct from ReLU and sigmoid networks (Fig. 2D). Furthermore, the embedding revealed that tanh RNNs with and without Dale’s connectivity constraint form a single cluster. In contrast, ReLU and sigmoid RNNs with and without Dale’s connectivity constraint form clusters which are clearly separable.

Thus, analyses of population trajectories and single-unit selectivity revealed that neural representations in tanh networks are distinct from those in ReLU and sigmoid RNNs. In addition, Dale’s connectivity constraint does not affect neural representations in tanh networks, contrasting with ReLU and sigmoid RNNs.

### Differences in dynamics across RNN architectures

Having observed distinct neural representations across RNN architectures, we next asked whether these differences may reflect distinct dynamical mechanisms used by the RNNs to solve the task. We characterized dynamical mechanisms by analyzing the fixed point configurations in RNNs with various architectures^13^. The fixed points are the points in the state space of a dynamical system where the flow field is equal to zero for a given constant input. Accordingly, once the RNN state is at a fixed point, it remains unchanged unless perturbed. Fixed point configurations are often used as a computationally tractable description of task-relevant dynamics in RNNs^5, 25^.

In each RNN, we computed the fixed points for each combination of task inputs (Methods: Fixed point finder). Specifically, for the CDDM task, we computed fixed points with both the context and sensory inputs held constant for a total of 50 distinct input combinations (5 relevant and 5 irrelevant coherences in 2 contexts). We aggregated the fixed points from all inputs and assessed their stability, categorizing each fixed point as either stable or unstable. We then compared the resulting sets of fixed points across RNNs with different architectures.

First, we visualized the fixed point configurations of example RNNs by projecting them on a plane spanned by the first two PCs (Fig. 3A). ReLU and sigmoid RNNs showed similar fixed point configurations. Their fixed points were clearly separated along the second PC according to the context cue. Within each context, the stable fixed points clustered at the far ends along the first PC, corresponding to left and right choices, with the unstable points located in between. The stable fixed points in ReLU and sigmoid RNNs formed elongated clusters, indicating that irrelevant stimulus is still represented, albeit to a limited degree. In contrast, fixed points of tanh RNNs form sheet-like structures, with irrelevant information being less suppressed, as evidenced by the nearly uniform distribution of fixed points across the sheets. While the fixed point configurations of tanh networks were unaffected by the Dale’s connectivity constraint, ReLU and sigmoid RNNs showed less variability in the fixed point configuration across networks under this constraint compared to when it was absent.

**Figure 3.**
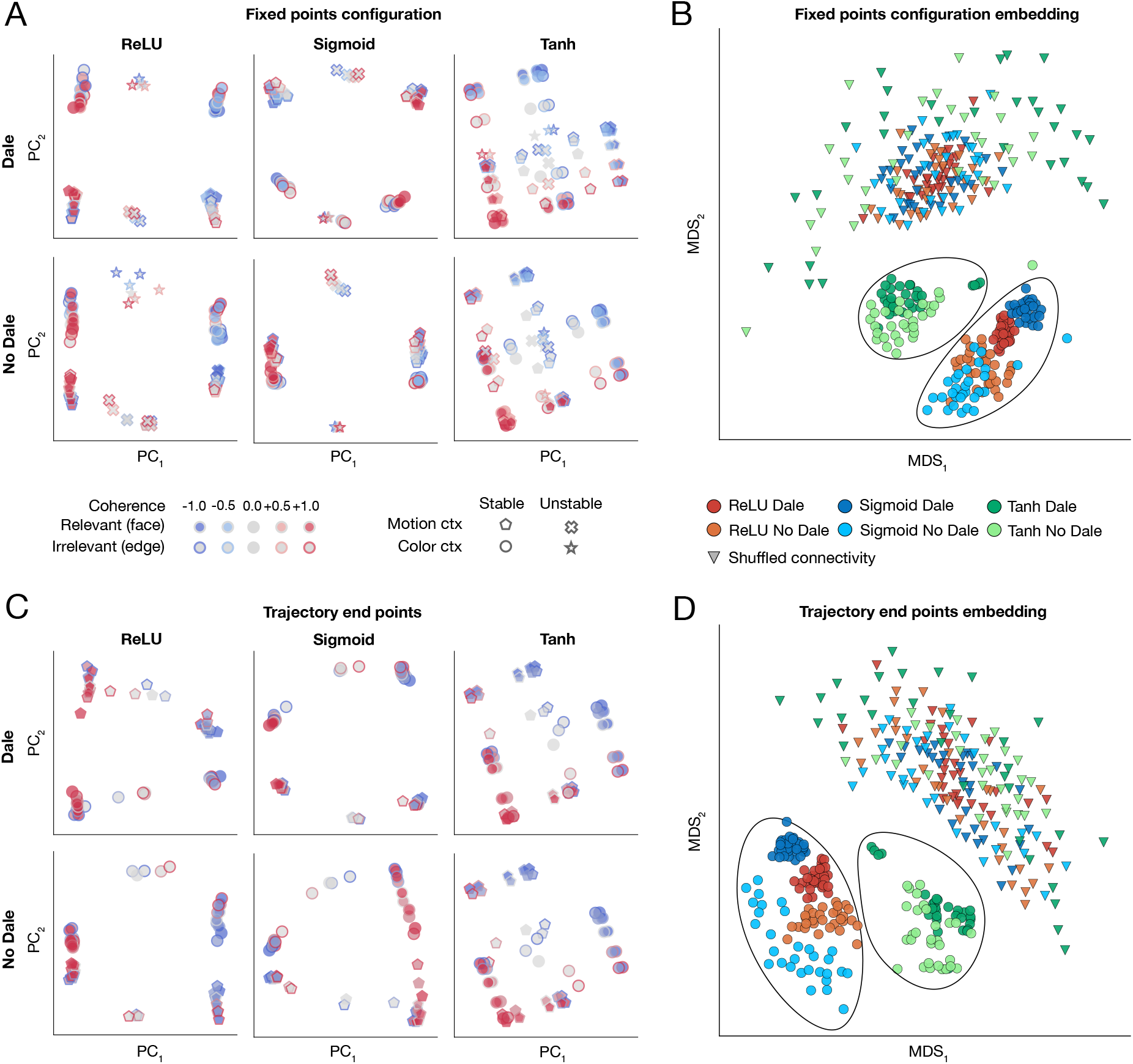
Comparison of fixed point and trajectory end point configurations across six RNN architectures trained on CDDM task. **A**. Fixed point configurations visualized by projecting onto the first two PCs in example RNNs with different activation functions (columns) and connectivity constraints (rows). We compute stable and unstable fixed points for each of 50 possible input combinations to the RNN (2 contexts cues, 5 relevant and 5 irrelevant coherences). **B**. MDS embedding of fixed point configurations across all RNNs. Each point represents fixed point configuration of one RNN. ReLU and sigmoid networks form clusters distinct from tanh networks. Triangles represent fixed points of the same RNNs with shuffled connectivity matrices, used as a control. **C**. Trajectory end point configurations visualized by projecting onto the first two PCs mirror the configurations of stable fixed points in example RNNs. **D**. MDS embedding of trajectory end point configurations across all RNNs.

To systematically quantify these differences across all RNNs, we embedded their fixed point configurations in a two-dimensional space using MDS (Fig. 3B, Methods: Analysis of fixed points). We computed distance between a pair of RNNs as MSE between the corresponding sets of fixed points aligned with a custom registration algorithm, which accounted for the fixed point type (stable or unstable) for each input. The resulting MDS embedding confirmed our initial observations: while all clusters corresponding to different architectures were separable, the tanh RNNs formed a cluster that was positioned further away from the others. We further verified these results by analyzing configurations of trajectory end points (network state at the last time step of a trial), which correlate with the arrangement of stable fixed points. As expected, the trajectory end point configurations mirrored the fixed point arrangement (Fig. 3C), and the MDS embedding of trajectory end points further reinforced our finding that tanh RNNs were distinct from both ReLU and sigmoid networks (Fig. 3D).

### RNN with varying architectures rely on different circuit mechanisms

We observed differences in population trajectories, single-unit selectivity, and fixed point configurations across RNNs with varying architectures. We next tested whether these variations in neural representations and dynamics reflect differences in the circuit solutions each RNN class discovers for the same task.

To identify the circuit mechanism used by each RNN to solve the CDDM task, we fit both its neural responses and task behavior with a latent circuit model^10^ (Methods: Latent circuit inference). Specifically, we fit RNN responses as a linear embedding of dynamics generated by a low-dimensional RNN—the latent circuit—with a matching activation function. We also require the latent circuit to reproduce task outputs. Thus, the latent circuit model infers a low-dimensional circuit mechanism generating task-relevant dynamics in the RNN. We inferred latent circuit mechanisms for top *n* = 10 RNNs with the best performance on CDDM task in each architectural class. All latent circuits produced accurate fits while also successfully solving the CDDM task (Table 2).

**Table 2.**
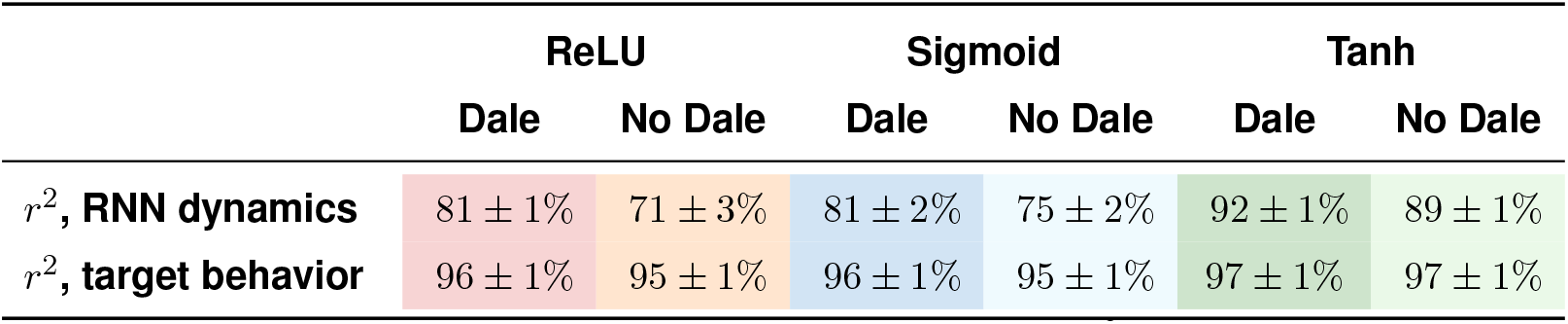
Latent circuit fit accuracy. Coefficient of determination *r*^2^ for the trajectories (upper row) and behavioral output (lower row) of the fitted latent circuit compared to the target RNN. The data are mean ±std across the best-fitting latent circuits for each of the top 10 RNNs trained on CDDM task within each architectural class.

|  | ReLU |  | Sigmoid |  | Tanh |  |
| --- | --- | --- | --- | --- | --- | --- |
|  | Dale | No Dale | Dale | No Dale | Dale | No Dale |
| $r^2$, **RNN dynamics** | $81 \pm 1\%$ | $71 \pm 3\%$ | $81 \pm 2\%$ | $75 \pm 2\%$ | $92 \pm 1\%$ | $89 \pm 1\%$ |
| $r^2$, **target behavior** | $96 \pm 1\%$ | $95 \pm 1\%$ | $96 \pm 1\%$ | $95 \pm 1\%$ | $97 \pm 1\%$ | $97 \pm 1\%$ |

The inferred latent circuit connectivity revealed that ReLU and sigmoid RNNs rely on a mechanism distinct from that of tanh RNNs to select relevant stimuli in the CDDM task. In ReLU and sigmoid RNNs, context nodes inhibit sensory nodes representing irrelevant stimuli in each context (Fig. 4A, e.g., motion context node inhibits sensory nodes representing color). Since the activity of irrelevant sensory nodes is suppressed, only the relevant nodes can affect the behavioral choice output^10^. In contrast, tanh RNNs block irrelevant stimuli using a qualitatively different circuit solution (Fig. 4A). In tanh RNNs, the active context node drives the nodes representing relevant and irrelevant stimuli to the opposite saturation regions of the tanh activation function. Prior to stimulus presentation, the RNN output is precisely zero due to a stalemate between the negatively saturated relevant nodes and positively saturated irrelevant nodes. A positive stimulus drives the relevant nodes to the steep region of the tanh activation function affecting the output, while stimulus input does not change the activity of the positively saturated irrelevant nodes and hence has no effect on choice.

**Figure 4.**
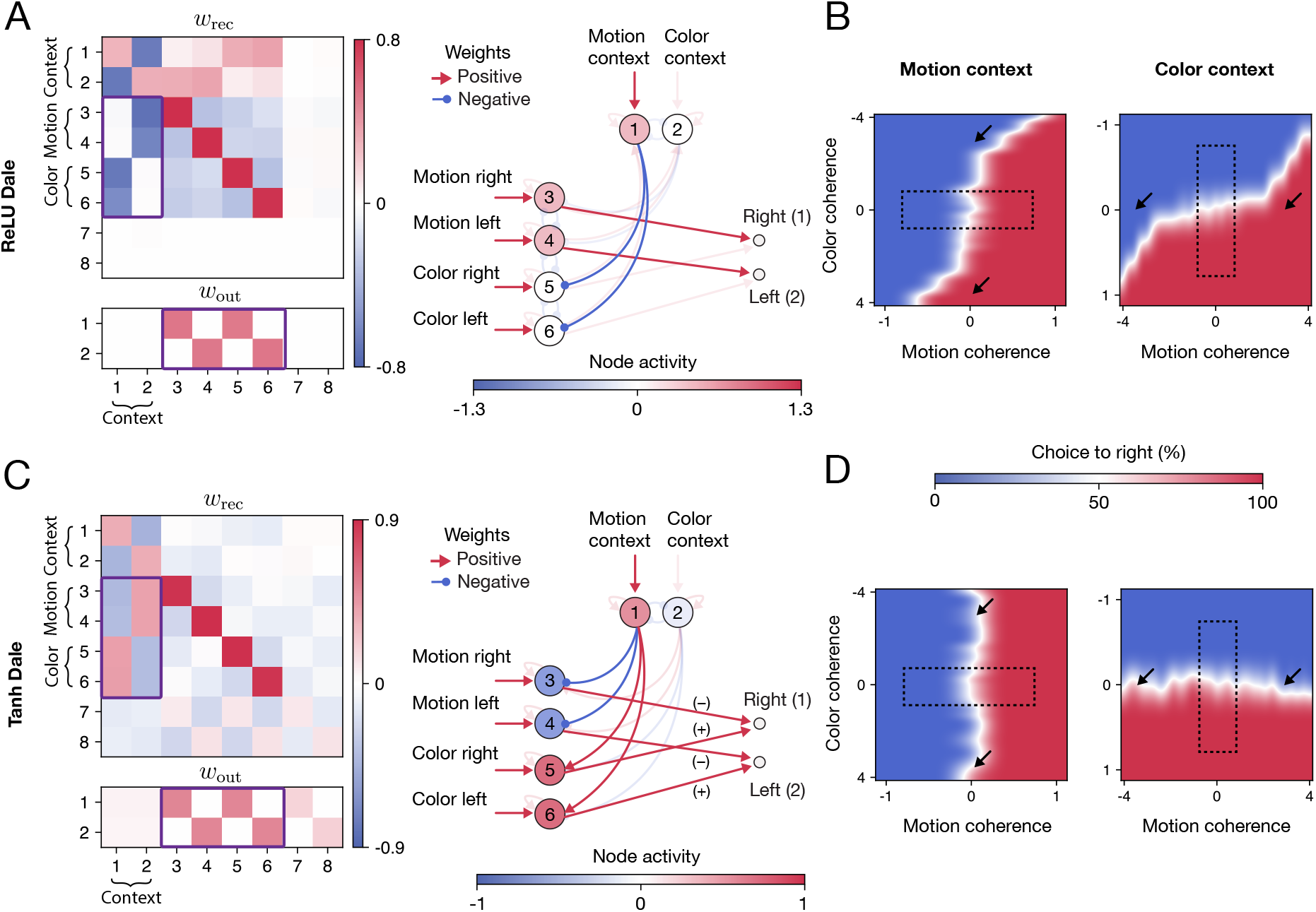
Distinct circuit solutions for CDDM task in ReLU versus tanh RNNs. **A**. Latent circuit connectivity inferred from responses of a ReLU RNN trained on the CDDM task, including recurrent (*w*_rec_) and output (*w*_out_) connectivity matrices (left). A simplified circuit diagram (right) highlights only the key nodes and connections for clarity, with the filled color representing the activity of each node on a motion context trial with both context cue and zero-coherence stimuli present. The latent circuit reveals a mechanism for selecting relevant stimuli based on inhibition of nodes representing irrelevant stimuli. Sensory nodes representing motion and color stimuli project to the corresponding outputs (purple rectangle in *w*_out_, red arrows in the circuit diagram). Inhibitory connections from the context to sensory nodes (purple rectangle in *w*_rec_, blue arrows in the circuit diagram) suppress the irrelevant stimulus representations in each context. **B**. Psychometric functions of the ReLU RNN for stimuli extending beyond the range used during training (rectangle indicates the stimulus range on which the RNN was trained). The network becomes sensitive to irrelevant stimuli with increased amplitude evident as a rotation of the decision boundary (arrows). Sigmoid RNNs showed qualitatively similar latent circuit mechanism and out-of-distribution behavior (data not shown). **C**. Same as A for latent circuit connectivity inferred from responses of a tanh RNN trained on the CDDM task. The latent connectivity reveals a mechanism for selecting relevant stimuli based on saturation of nodes representing irrelevant stimuli. Context nodes drive the nodes receiving irrelevant stimuli into the positive saturation region of the tanh activation function, while pushing the nodes receiving relevant stimuli into the negative saturation region (purple rectangle in *w*_rec_). Prior to stimulus presentation, the negative activity of relevant nodes and positive activity of irrelevant nodes cancel each other at the output (purple rectangle in *w*_out_; red arrows in the circuit diagram). Relevant stimuli drive the relevant nodes into the steep region of the tanh activation function, allowing them to affect the output. At the same time, irrelevant stimuli drive the irrelevant nodes further into the saturation region, where their activity remains unchanged, thus having no effect on the output. **D**. Same as B for the tanh RNN. Strong irrelevant stimuli do not affect the network’s choice (arrows).

These different circuit mechanisms make distinct predictions for how networks will respond to out-of-distribution inputs. The tanh circuit predicts that increasing the amplitude of the irrelevant stimulus beyond the range used during training will not affect the behavioral output, because it will push the irrelevant nodes further in the positive tanh saturation region without changing their activity. In contrast, the ReLU circuit predicts that a sufficiently strong irrelevant input will overcome the inhibition of the irrelevant nodes by the context nodes, thereby allowing the irrelevant stimulus to bias the behavioral output. These predictions were clearly borne out by data. When exposed to irrelevant inputs with amplitudes greater than during training, the ReLU and sigmoid RNNs become sensitive to these irrelevant stimuli, evident as a rotation of the decision boundary in the psychometric function (Fig. 4B). In contrast, strong irrelevant stimuli with amplitudes beyond the training range have no effect on the psychometric function of tanh networks (Fig. 4D). These results demonstrate that circuit mechanisms define how networks generalize to out-of-distribution inputs and that different RNN architectures have inductive biases favoring different circuit mechanisms.

Together, our results show that differences in population trajectories, single-unit selectivity, and fixed point configurations can indicate distinct circuit solutions discovered by RNNs for the same task. Moreover, RNN architectures impose inductive biases that favor specific circuit solutions, highlighting the importance of the architectural choice in modeling biological data.

### Differences in RNN architectures manifest across tasks

Are differences in neural representations and dynamics across RNN architectures specific to the CDDM task, or do they appear in other tasks as well? To answer this question, we trained ensembles of RNNs with all six architectures to perform the Go/NoGo and Memory Number tasks (Fig. 1B,C, Methods: Go/NoGo and Memory Number tasks). We compared neural representations and dynamics in these RNNs using population trajectories, singleunit selectivity, and fixed point configurations. In these both tasks, example ReLU and sigmoid RNNs showed qualitatively similar projected trajectories, whereas the trajectories of tanh networks significantly differed from that of ReLU and sigmoid RNNs (Fig. 5A,B). Specifically, low-amplitude inputs generated highly compressed trajectories as compared to large excursions through the state space produced by large inputs in ReLU and sigmoid RNNs. In contrast, the extent of trajectories was more similar between high- and low-amplitude inputs in tanh RNNs.

**Figure 5.**
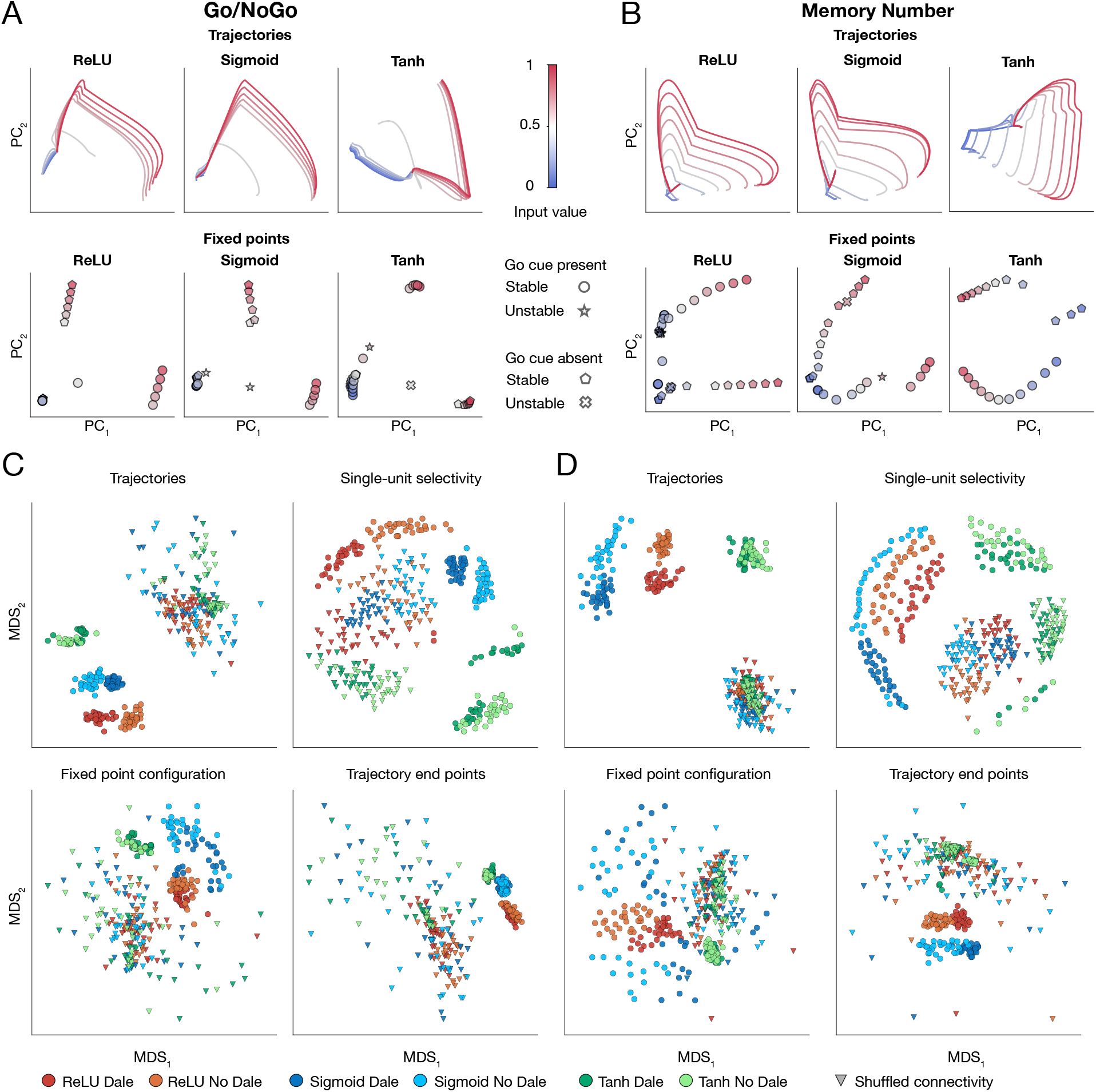
Comparison of population trajectories, single-unit selectivity, fixed points, and trajectory end point configurations across RNN architectures for Go/NoGo and Memory Number tasks. **A**. Population trajectories (upper row) and fixed point configurations (lower row) projected onto the first two PCs for Go/NoGo task for example ReLU, sigmoid and tanh RNNs with Dale connectivity constraint. **B**. Same as A for Memory Number task. **C**. MDS embeddings of population trajectories, single-unit selectivity, fixed points and trajectory end points across all RNNs trained on Go/NoGo task. Each point in the embedding space represents a single RNN. Triangles represent trajectories of the same RNNs with shuffled connectivity matrices, used as a control. In all embedding spaces, tanh networks form distinctly separated clusters, while ReLU and sigmoid networks, although distinct, are typically closer to each another. **D**. Same as C for Memory Number task.

Visualizing the distribution of each metric across all RNNs using MDS embedding, we observe that ReLU, sigmoid, and tanh networks form separate clusters for both tasks (Fig. 5C,D). Although ReLU and sigmoid networks form distinct clusters in each embedding space, they are closer to each other than to tanh RNNs, confirming the consistent idiosyncrasy of tanh networks across all three tasks. In the MDS embeddings of all metrics, Dale and No Dale networks with the same activation function formed partially overlapping yet distinct subclusters, positioned closer to each other than to any subclusters corresponding to other activation functions. Thus, while the constraint imposed by Dale’s law had some influence, the choice of activation function had significantly greater effect on representations and dynamics in RNNs. These results further support the conclusion that the choice of activation function impacts the emergent task solutions in RNNs, with tanh networks standing out as the most distinct from ReLU and sigmoid counterparts.

## Discussion

We show that RNN architectures confer inductive biases that influence neural population dynamics, single-unit selectivity, and circuit mechanisms emerging through training on cognitive tasks. Different circuit mechanisms manifest in diverging behavior on out-of-distribution inputs, demonstrating that these differences reflect fundamentally distinct task solutions rather than mere trivial variations. Task-optimized RNNs are widely used to generate hypotheses for how the brain may solve cognitive tasks, yet the choice of activation function is often tacitly assumed inconsequential for the resulting mechanisms, reflected in a variety of activation functions used across modeling studies^3–6, 10, 14–21^. Our findings indicate that different architectures can yield disparate circuit solutions, which may vary substantially in how well they align with the circuit mechanisms in the brain. RNNs can also be directly optimized to reproduce neural recording data, such that each RNN unit tracks the activity of one experimental neuron^7–9, 26^. Our results have broader implications for this approach as well, as some architectures may be more amenable than others to replicating neural recordings. In addition, it remains an open question whether two architectures that can equally well fit the same neural responses converge on the same circuit solution. Therefore, the choice of RNN architecture cannot be ignored when modeling biological systems, as these choices may bias the inferred solutions and their relevance to neural processes.

While we trained six architectures on three tasks to achieve comparable performance, some architectures may be better suited for specific tasks than others^27^, much like how specific basis functions are more effective at approximating certain types of functions. In the brain, the suitability of different single-unit dynamics for different information processing needs may be supported by a diverse array of cell types, each potentially fine-tuned for specific computations^28–30^. Our Dale-constrained ReLU and sigmoid networks incorporate two basic cell types: excitatory and inhibitory units. Neural representations in these RNNs with two cell types differed from their counterparts without Dale’s constrain, suggesting that the existence of multiple cell types also affects task solutions emerging in RNNs. Moreover, incorporating multiple cell types with diverse activation functions improved the image classification accuracy of convolutional neural networks compared to conventional homogeneous architecture^31, 32^. Thus, equipping RNN models with multiple unit types, featuring different activation functions and connectivity constraints that correspond to biological cell types, may result in closer alignment of RNNs with biological circuits and higher computational efficiency.

Some activation functions may be effective for solving tasks in artificial networks, but not correspond to feasible single-unit dynamics in biological networks. For example, biological neurons cannot produce negative firing rates, making tanh units significantly less biological plausible. Tanh units reverse the sign of their synaptic effect depending on their activity state, a feature that can be useful for solving certain tasks but not observed in biological neurons. In tanh networks, Dale’s connectivity constraint loses its biological relevance in defining excitatory and inhibitory cell types. Consistently, we observed that this constraint does not influence neural representations in tanh RNNs. With this distinct singleunit property, tanh RNNs consistently diverged from the more biologically plausible ReLU and sigmoid RNNs across all metrics and tasks we examined.

Moreover, tanh RNNs produced a circuit mechanism for context-dependent decision-making that allows them to block irrelevant stimuli even with arbitrary large amplitudes. In contrast, ReLU and sigmoid networks become increasingly sensitive to stronger irrelevant stimuli. In practice, human behavior is often influenced by strong irrelevant stimuli, as demonstrated by the Stroop effect, where error rates increase when responding to incongruent stimuli^33^. Thus, although tanh RNNs discover a more robust solution to this task, the ReLU and sigmoid networks exhibit behavior that aligns more closely with experimentally observed psychophysical patterns. These observations suggest that biological implausibility of single-unit dynamics may translate into circuit mechanisms and behavior that deviate from biological systems.

We observed that more biologically plausible ReLU and sigmoid RNNs show less heterogeneous single-unit selectivity under Dale’s connectivity constraint compared to their unconstrained counterparts (Fig. 2C). Consistently, latent circuits of the same dimensionality captured more variance in RNNs with Dale’s connectivity constraint than in those without it (Table 2), facilitating analysis of these networks. Dale’s connectivity constrain introduces only a coarse level of biological realism, which may not offer a sufficient inductive bias to capture complex circuit solutions implemented in biological networks. Additional biological constrains may gradually steer artificial models to better align with biological neural circuits, enhancing their ability to generate more relevant hypotheses.

Although RNNs are coarse models of biological neural networks and their units’ activation functions oversimplify single-neuron dynamics, our findings emphasize the critical role of the RNN architecture in generating biologically relevant hypotheses. RNN architectures introduce inductive biases for the emergent task solutions, which can lead to distinct hypotheses about how the brain solves these tasks. While the question of which RNN architectures best align with biological data remains open, our results suggest that tanh activation function may not be the optimal choice. Directly comparing neural recording data with the representations and dynamics of RNNs across different architectures will help determine which architecture best suits the modeling of biological neural networks.

## Methods

### RNN architectures and training procedure

For each of the six architectures ({Dale, No Dale} ⊗ {ReLU, sigmoid, tanh}), we trained 100 fully connected RNNs, each with *N* = 100 units, to solve cognitive tasks. The RNN dynamics are described by the equations:

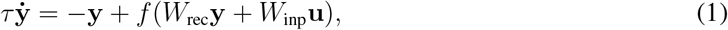

where *f* is the activation function (ReLU, sigmoid or tanh). The sigmoid function is defined as sigmoid(**x**) = 1*/*(1 + *e*^−7.5**x**^) with the slope 7.5. The ReLU function is defined as ReLU(**x**) = max(0, **x**), and tanh function is tanh(**x**) = (*e*^**x**^ − *e*^−**x**^)*/*(*e*^**x**^ + *e*^−**x**^).

The RNNs are trained by minimizing the loss function:

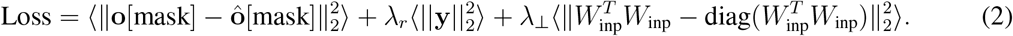

Here **o** are the target outputs and **ô**= *W*_out_**y** are the RNN outputs, both with dimensionality of (*n*_output_, *T, K*), where *T* is the number of time steps in the trial and *K* is the number of trials, and ⟨·⟩ denotes the mean over all dimensions. The indexing matrix `mask` specifies the time steps on which the loss is evaluated for each task. The regularization term 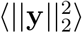 biases the training procedure to find solutions with moderate activity levels. The last term in the loss function enforces orthogonality between the columns of *W*_inp_, which empirically improves the convergence of RNN training. In addition, some units may become inactive during training, so that they do not contribute to the computations. We empirically found that introducing the orthogonality term helped to decrease the number of silent neurons as well. We initialize the RNN connectivity matrix as described previously^3^ and optimize the loss function using Adam optimizer in pytorch, with default hyperparameters: learning rate *α* = 0.001, *β*_1_ = 0.9 *β*_2_ = 0.999, *ϵ* = 10^−8^.

The RNN output was obtained by running the RNN’s dynamics forward for a given batch of inputs. We discretize the RNN dynamics using the first-order Euler scheme with a time-step *dt* = 1 ms and add a noise term in the discretized equation to obtain:

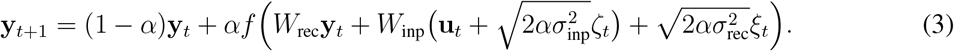

Here *α* = *dt/τ*, and *ξ* and ζ are vectors of random variables, each element of which is sampled from the standard normal distribution *N* (0, 1). The hyperparameters used for RNN training are provided in Table 3. RNNs were trained on the CDDM and Go/NoGo tasks with *λ*_*r*_ = 0.5 for *n*_iter_ = 5, 000 iterations. RNNs were trained on the Memory Number task first with *λ*_*r*_ = 0 for *n*_iter_ = 6, 000 and then with *λ*_*r*_ = 0.3 for additional *n*_iter_ = 6, 000.

**Table 3.**
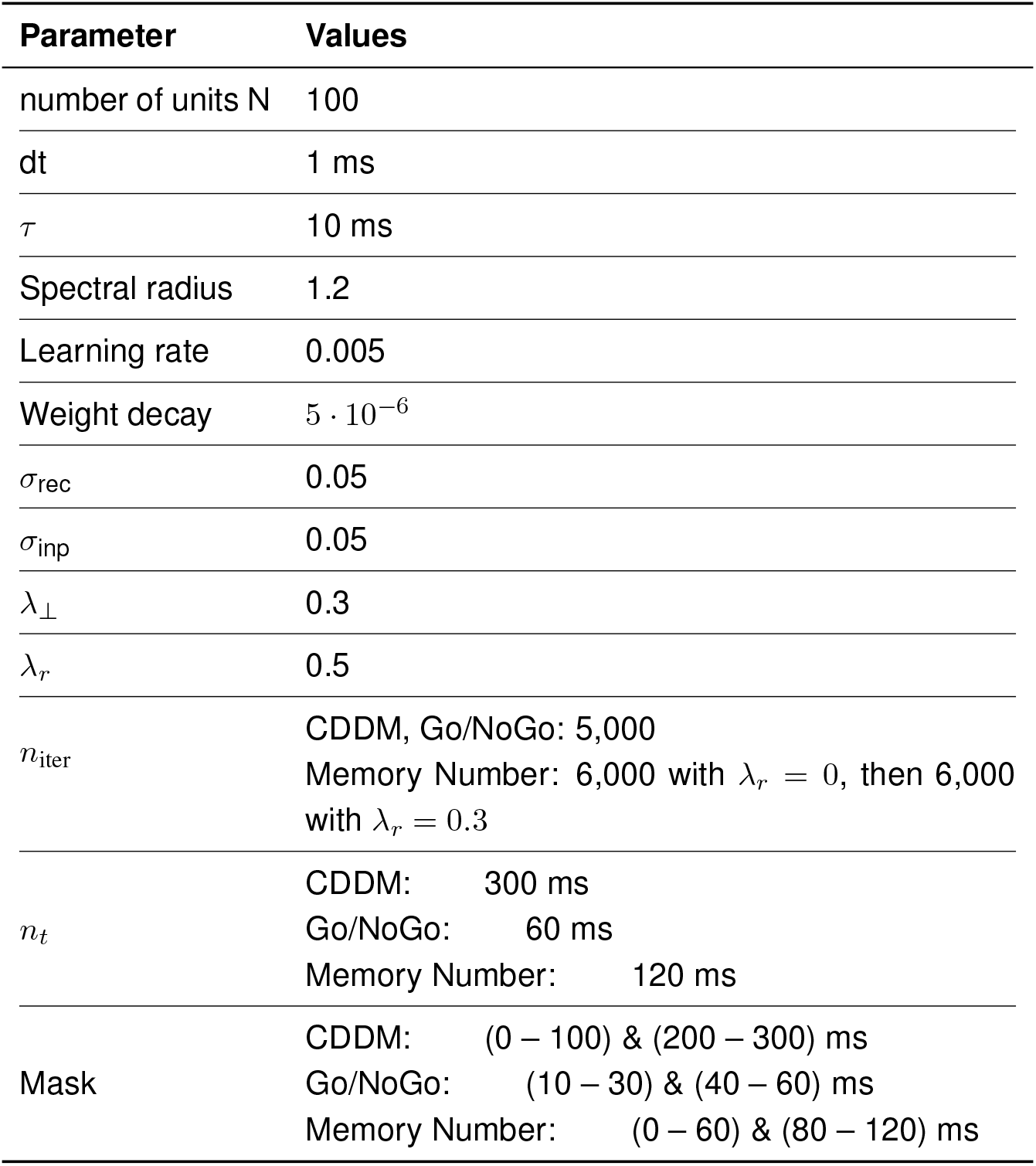
RNN training hyperparameters.

| Parameter | Values |
| --- | --- |
| number of units N | 100 |
| dt | 1 ms |
| $\tau$ | 10 ms |
| Spectral radius | 1.2 |
| Learning rate | 0.005 |
| Weight decay | $5 \cdot 10^{-6}$ |
| $\sigma_{\text{rec}}$ | 0.05 |
| $\sigma_{\text{inp}}$ | 0.05 |
| $\lambda_{\perp}$ | 0.3 |
| $\lambda_r$ | 0.5 |
| $n_{\text{iter}}$ | CDDM, Go/NoGo: 5,000<br>Memory Number: 6,000 with $\lambda_r = 0$, then 6,000 with $\lambda_r = 0.3$ |
| $n_t$ | CDDM: 300 ms<br>Go/NoGo: 60 ms<br>Memory Number: 120 ms |
| Mask | CDDM: (0 – 100) & (200 – 300) ms<br>Go/NoGo: (10 – 30) & (40 – 60) ms<br>Memory Number: (0 – 60) & (80 – 120) ms |

### Context-dependent decision-making task

The task structure is presented in Fig. 1A. Two mutually exclusive context channels signal either ‘Motion’ or ‘Color’ context. For a given context, a constant input with an amplitude of 1 is supplied through the corresponding channel for the entire trial duration. Sensory stimuli with two modalities (‘Motion’ and ‘Color’) are each supplied through two corresponding input channels, encoding momentary evidence for choosing either the right or left response. Within each sensory modality, the mean difference between inputs on two channels represents the stimulus coherence, with values ranging from −1 to +1. During training, we used a discrete set of 15 coherences for each sensory modality: *c* = {0, ±0.01, ±0.03, ±0.06, ±0.13, ±0.25, ±0.5, ±1}. The coherence *c* was translated into two sensory inputs as [(1 + *c*)*/*2, (1−*c*)*/*2]. For 300 time steps on a trial, the (6, 300)dimensional input-stream array was calculated based on the triplet (binary context, motion coherence, color coherence), generating *N*_batch_ = 2 × 15 × 15 = 450 distinct trial conditions.

On each trial, the target output was set to 0 for each time step *t <* 100 ms. During the decision period *t >* 200 ms, the target was set as follows: if the relevant coherence (e.g., coherence of ‘Motion’ stimuli on a ‘Motion’ context trial) was positive, the target for ‘Right’ output channel was set to 1 from 200 ms onward. If the relevant coherence was negative, the target for ‘Left’ output channel was set to 1 instead. If the relevant coherence was 0, both output targets were set to 0. The target was specified for only a subset of time steps, forming a training mask: (0 − 100) & (100 − 200) ms, allowing the network to integrate the decision without penalty in the intervening period.

### Go/NoGo and Memory Number tasks

The structure of these tasks is presented in Fig. 1B,C. For both tasks, we used 11 uniformly spaced input values ℐ, ranging from 0 to 1, delivered through the first input channel. The ‘Go Cue’ input is delivered through the second channel and activated only at time *t*_GoCue_ at the end of the trial, signaling that the RNN is required to respond. Finally, a constant bias input with amplitude of 1 is supplied via the third channel throughout the entire trial duration. In the Go/NoGo task, the input value ℐ was provided for the entire trial duration of 60 ms. The target output is determined as 0 before, and Θ(ℐ − 0.5) after the Go Cue onset, where Θ is the Heaviside step function (Fig. 1B). If the input value was exactly 0.5, the network was required to output 0.5 after the Go Cue. In the Memory Number task, the input value ℐ was present only for 10 ms, with the randomized stimulus onset window *t*_stim_ ∼ *U* (0, 20) ms (Fig. 1C). The target output value was set to be 0 before the Go Cue and the input value ℐ afterwards. The onset of the Go Cue was set to *t*_GoCue_ = 30 ms for the Go/NoGo task, and *t*_GoCue_ = 70 ms for the Memory Number task.

### Controls: RNNs with shuffled connectivity

For each of the analyzed RNNs, we produced another RNN with shuffled connectivity as a control. To shuffle the connectivity, we randomly permute each row *R*_*i*_ in the input matrix *W*_inp_ (*i*^*th*^ row contains all inputs to unit *i*). We also randomly permute nondiagonal elements of each column in the recurrent matrix *W*_rec_ (*i*^*th*^ column contains all outputs of unit *i*). We keep the diagonal elements in *W*_rec_ unchanged to preserve self-excitation of each unit.

### Analysis of population trajectories

We analyzed 30 RNNs with the best task performance from each architecture. We simulated each RNN (including the corresponding control RNNs) to acquire a tensor of neural responses *Z* with dimensionality (*N, T, K*), where *N* is the number of units in the network, *T* is the number of time steps in a trial, and *K* is the number of trials. We reshape the neural response tensor *Z* to obtain a matrix *X* with dimensionality (*N, T* · *K*). We then obtain a denoised matrix *F* with dimensionality (*n*_PC_, *T* · *K*) by projecting matrix *X* onto the first *n*_PC_ = 10 PCs along the *first* dimension, capturing ⩾ 93% of variance in each instance across all RNNs and tasks. Reshaping matrix *F* back into a three-dimensional tensor, we obtain a denoised tensor 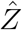 with dimensionality (*n*_PC_, *T, K*) containing reduced population trajectories. We further normalized the reduced trajectory tensor 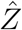 by its variance, so that the reduced trajectory tensors have the same scale across all RNNs.

To obtain an MDS embedding of the reduced trajectories, we compute a distance matrix between reduced trajectory tensors 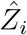 and 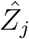 for each pair of RNNs *i* and *j*. First, we obtain the optimal linear transformation between the matrices *F*_*i*_ and *F*_*j*_ corresponding to 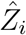 and 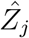 using linear least squares regression with the function `numpy.linalg.lstsq` in python. We perform two regression analyses: first regressing *F*_*i*_ onto *F*_*j*_ and then *F*_*j*_ onto *F*_*i*_, resulting in two linear transformations *M*_*ij*_ and *M*_*ji*_ and two scores, score_1_ = ∥*F*_*i*_*M*_*ij*_ − *F*_*j*_∥_2_ and score_2_ = ∥*F*_*j*_*M*_*ji*_ − *F*_*i*_∥_2_. We then compute the distance between two trajectory-tensors as the average of two scores: *d*_*ij*_ = *d*_*ji*_ = (score_1_ + score_2_)*/*2. We use these pairwise distances to compute MDS embedding with the function `sklearn.manifold.MDS` from `sklearn` package in python.

### Analysis of single-unit selectivity

For each RNN (including the control RNNs), we start with the same neural response tensor *Z* as for the analysis of population trajectories. We reshape *Z* to obtain matrix *X* with dimensionality (*N, T* · *K*). We then obtain a denoised matrix *G* with dimensionality (*N, n*_PC_) by projecting matrix *X* onto the first *n*_PC_ = 10 PCs along the *second* dimension, capturing above 90% of variance in each instance across all RNNs and tasks. We further normalize the resulting single-unit selectivity matrix *G* by its variance, so that single-unit selectivity matrices have the same scale across all RNNs.

To obtain an MDS embedding, we compute a distance matrix between the single-unit selectivity matrices *G*_*i*_ and *G*_*j*_ for each pair of RNNs *i* and *j*. To compute the distance between *G*_*i*_ and *G*_*j*_, we view each RNN unit as a point in *n*_PC_-dimensional selectivity space. We then register the point configurations of two RNNs with an optimal orthogonal transformation that allows for one-to-many mapping. To register the points, we use iterative closest point (ICP) registration algorithm (Methods: Iterative closest point registration), since there is no one-to-one correspondence between units in two RNNs. We perform the ICP registration two times, registering *G*_*i*_ to *G*_*j*_ and then *G*_*j*_ to *G*_*i*_, producing score_1_ and score_2_. We then set the distances *d*_*ij*_ = *d*_*ji*_ = (score_1_ + score_2_)*/*2.

### Fixed point finder

To find fixed points of an RNN, we use a custom fixed point finder algorithm. For each constant input **u**, we search for fixed points by minimizing the right hand side in Eq. 1, *F* (**y, u**) = −**y** + *f* (*W*_rec_**y** + *W*_inp_**u**) with `scipy.optimize.fsolve` function from `scipy.optimize` package in python. We accept point **y**^∗^ as a fixed point if 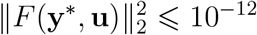.The `fsolve` function also takes the Jacobian matrix *J*(**y, u**) = ∂*F* (**y, u**)*/*∂**y** of the RNN as an additional argument to enhance the efficiency of the optimization process. We initialize the minimization at a value **y**_0_ sampled randomly from the RNN trajectories: we choose a random trajectory *k* from *K* trials, and then a random time-step *t* from the interval (*n*_*t*_*/*2, *n*_*t*_), that is, from the second half of the trial. We then add Gaussian noise *ξ* ∼ *N* (0, 0.01) to each coordinate of the sampled point to obtain the initial condition **y**_0_.

To find multiple fixed points for the same input **u**, we search for fixed points starting from multiple initial conditions within an iterative loop. On each iteration of this loop, we sample a new initial condition and perform the minimization to find a fixed point. We then compare this newly found fixed point 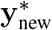 to all previously found fixed points 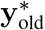.If the distance 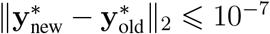,then we discard the new fixed point because it lies too close to one of the previously found fixed points. This iterative loop continues until either 100 distinct fixed points were found in total or no new fixed points were found for 100 consecutive iterations.

We determine the fixed point type (stable or unstable) by computing the principal eigenvalue *λ*_0_ of the Jacobian *J*(**y, u**) evaluated at the fixed point. We classify the fixed point as stable if ℝe(*λ*_0_) ⩽ 0 and otherwise as unstable.

### Analysis of fixed points

For each RNN (including the control RNNs), we computed fixed points for each combination of input stimuli using a custom fixed point finder algorithm (Methods: Fixed point finder), obtaining a fixed point configuration, which is a set of stable and unstable fixed points for different combinations of inputs. We collect the coordinates of all fixed points in a matrix *P* with dimensions (*N*_*p*_, *N*), where *N*_*p*_ is the total number of fixed points (both stable and unstable) across all the inputs and *N* is the number of units. We reduce the second dimension of the matrix *P* by projecting the fixed points onto the first *n*_PC_ = 7 principal components. We further normalized the resulting matrix by its variance, so that these fixed point configurations have the same scale across all RNNs, obtaining a matrix 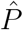 for each RNN. Throughout the transformations, we keep each fixed point tagged by its type and the corresponding input for which it was computed.

To obtain an MDS embedding, we compute a distance matrix between fixed point configurations 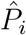 and 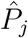 for each pair of RNNs *i* and *j*. To compute the distances between the two projected fixed point configurations 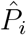 and 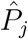,we compute an optimal orthogonal transformation between the two sets of projected fixed points using orthogonal Procrustes with iterative closest point registration (Methods: Iterative closest point registration). When matching the projected fixed points, we restricted matches to the fixed points with the same tag (of the same type and obtained for the same input). We perform the ICP registration two times, registering 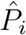 to 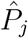 and then 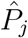 to 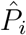,resulting in two scores score_1_ and score_2_. We then set the distances *d*_*ij*_ = *d*_*ji*_ = (score_1_ + score_2_)*/*2. Using the distance matrix, we then obtain MDS embedding.

### Analysis of trajectory end point configurations

For each RNN (including the control RNNs), we use the same neural response tensor *Z* as for the analysis of population trajectories. We then restrict the data to the last time step of each trial, resulting in (*K, N*) dimensional matrix *S* for each RNN containing the trajectory end point configuration. We further project the trial end points in *S* onto first *n*_PC_ = 10 principal components, obtaining (*K, n*_PC_)-dimensional matrix Ŝ. Finally, we normalize each trajectory end point configuration matrix Ŝ by its variance, so that these end point configurations have the same scale across all RNNs. We compute the distance between two matrices Ŝ_*i*_ and Ŝ_*j*_ for RNNs *i* and *j* using the same procedure as for the population trajectories matrices *F* (Methods: Analysis of population trajectories). Using the distance matrix, we then obtain MDS embedding.

### Iterative closest point registration

To register the point clouds (Methods: Single-unit selectivity and Methods: Fixed point analysis), we use an iterative closest point (ICP) algorithm, which proceeds in four steps:

1. **Initialization**: Define a random orthogonal matrix *A* that transforms each point of the source point cloud *P*_source_ into *P*_source_*A*.
2. **Point Matching**: For each point in the target point cloud *P*_target_, find the closest point in the transformed source point cloud *P*_source_*A*. Construct a new matrix 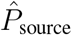 where the *i*^*th*^ point is the point from *P*_source_*A* closest to the *i*^*th*^ point in *P*_target_ (points in 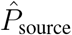 may repeat).
3. **Transformation Update**: Update the transformation matrix *A* to minimize the distance betweenv 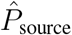 and *P*_target_ using orthogonal Procrustes method.
4. **Iteration**: Repeat steps 2 and 3 until convergence.

This algorithm iteratively refines the transformation to achieve optimal alignment between the source and target point clouds. Since this optimization is non-convex, it may converge to a local optimum. Therefore, we perform each optimization for *n*_tries_ = 31 starting with random initializations and keep the solution with the minimal mean squared error as a distance between the source and target point clouds.

To compute distances between the fixed point configurations (Methods: Analysis of fixed points), we modify the Point Matching step by restricting possible matches only to the points obtained for the same inputs and of the same type (stable or unstable).

### Latent circuit inference

To identify the circuit mechanism supporting the CDDM task execution in an RNN, we fit its responses and task behavior with the latent circuit model^10^. We model RNN responses *y* as a linear embedding of dynamics *x* generated by a low-dimensional RNN:

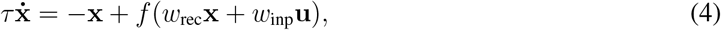

which we refer to as the latent circuit. Here *f* is the activation function matching the activation function of the RNN. We also require the latent circuit to reproduce task behavior via the output connectivity *w*_out_*x*.

To fit the latent circuit model, we first sample RNN trajectories *Z*, forming a (*N, T, K*)-dimensional tensor. We then reduce the dimensionality of *Z* using PCA to *N*_PC_ = 30, resulting in a tensor **z** with (*N*_PC_, *T, K*) dimensions, capturing above 99% of variance in *Z* for all RNNs we analyzed. We then infer the latent circuit parameters *w*_rec_, *w*_inp_, *w*_out_ and an orthonormal embedding matrix *Q* by minimizing the loss function:

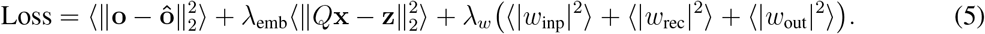

Here ⟨·⟩ denotes the mean over all dimensions of a tensor. Tensor **x** has the dimensionality (*n, T, K*), where *n* is the number of nodes in the latent circuit. This tensor **x** contains the activity of the latent circuit across *K* trials and *T* time steps per trial, and **y** is the corresponding activity tensor for the RNN. The (*N*_*PCs*_, *n*) dimensional orthonormal matrix *Q* embeds trajectories of the latent circuit **x** to match the RNN activity **z**, such that *z* ≈ *Qx*. Finally, **o** is the target circuit output, and the **ô** = *w*_out_*x* is the output produced by the latent circuit.

During optimization, we constrain the input matrix such that each input channel is connected to at most one latent node. To this end, we apply to the input matrix a mask, in which 1 indicates that the weight is allowed to change during training, and 0 indicates that the weight is fixed at 0. We design the mask such that each column has a single 1. Additionally, we constrain the elements of *w*_inp_ and *w*_out_ matrices to be non-negative.

We fitted latent circuit models to the 10 RNNs with the best CDDM task performance from each architecture. For each RNN, we fit 8-node latent circuit model ⩾ 30 times starting with random initializations, and take the best-fitting circuit as a converged solution. The hyperparameters for the latent circuit fitting are provided in Table 4.

**Table 4.** Hyperparameters for fitting latent circuit model.

| Parameter | Values |
| --- | --- |
| $N_{\text{PC}}$ | 30 |
| dt | 1ms |
| $\tau$ | 10 ms |
| $n_{\text{iter}}$ | 2,000 |
| Learning rate | 0.02 |
| $\lambda_w$ | 0.08 |

## Acknowledgements

This work was supported by National Institute of Health (NIH) grant RF1DA055666 (T.A.E.).

## Author contributions

P.T. and T.A.E. designed the research and developed the computational analysis framework. P.T. developed the code, performed computer simulations, and analyzed data. P.T. and T.A.E wrote the paper.

### Data availability

The synthetic data used in this study can be reproduced using the source code.

### Code availability

The code for RNN training is available as `trainRNNbrain` package on GitHub at https://github.com/engellab/trainRNNbrain. The code for latent circuit fitting is available on GitHub at https://github.com/engellab/latentcircuitinference. The code for the analysis of the RNNs and the relevant datasets are available on GitHub at https://github.com/engellab/ActivationMattersRNN.

